# Beyond dichotomy: diversity of ultrasonic vocalizations in rats

**DOI:** 10.1101/2024.12.09.627446

**Authors:** Reo Wada, Shiomi Hakataya, Ryosuke O. Tachibana, Tomoyo Isoguchi Shiramatsu, Tetsufumi Ito, Kouta Kanno, Riseru Koshiishi, Jumpei Matsumoto, Yumi Saito, Genta Toya, Shota Okabe, Kazuo Okanoya

## Abstract

Rats emit ultrasonic vocalizations (USVs) in diverse contexts, and these USVs are thought to serve many biological functions, such as socio-coordinating and alarming functions. In particular, rat USVs have been established as a good model for studying emotional expressions. Since the discovery of USVs in rats, it has gradually been revealed that two families of juvenile and adult rat USVs with distinct acoustic features are closely linked to different affective states: 50-kHz (high-short) calls are associated with positive affective states, whereas 22-kHz (low-long) calls are associated with negative affective states. Although various subtypes within high-short and low-long calls have been proposed and used, most studies adopt the dichotomous framework of first dividing USVs into high-short or low-long calls. The diversity of rat USVs may have been overlooked due to this high-short / low-long dichotomous framework, as such labeling could discourage reporting of actual measured acoustic properties. Considering that several recent studies claim to have found new USV categories outside this framework, we conducted a systematic survey of descriptions of rat USVs in the literature. We identified 15 articles reporting USVs that are outside the dichotomous framework. Our results support the existence of diverse USVs beyond the dichotomy, highlighting the importance of research on a broader range of vocalizations that might reflect complex affective states beyond the simple distinction between positive and negative.

## Introduction

Understanding animal emotions is of great interest to researchers across various disciplines, ranging from neuroscience to animal welfare. Rats (*Rattus norvegicus*), in particular, provide a good model for investigating emotional expression through their ultrasonic vocalizations (USVs), as these vocalizations are widely studied as indicators of emotional states [1]. Since the initial discovery of rat USVs [2], it has become increasingly evident that adult rats produce two distinct families of USVs, each linked to specific emotional contexts [3,4].

One family consists of calls with relatively high frequency and short duration (45–80 kHz, 5– 200 ms), traditionally labeled as “50-kHz USVs.” These calls are typically emitted during appetitive and positive contexts, including rough-and-tumble play [5], mating [6], tickling [7], social contact [8], and in response to the presentation or anticipation of rewarding stimuli such as amphetamine [9], cocaine [10], electric stimulation of the reward system in the brain [11], although they can also occur during aggression [12]. Another family is composed of calls with relatively low frequency and long duration (18–28 kHz, 500–2000 ms), traditionally labeled as “22-kHz USVs.” They tend to be produced in negative situations, including social defeat [13], exposure to electric footshocks [14], predator exposure [15], and in response to cues associated with aversive experiences, such as withdrawal from cocaine and opiates [16]. In the present study, we refer to the two families as “high-short (HS)” and “low-long (LL)” USVs (***Fig. 1A***), which helps avoid confusion with actual frequency values reported in the literature.

**Figure 1.**
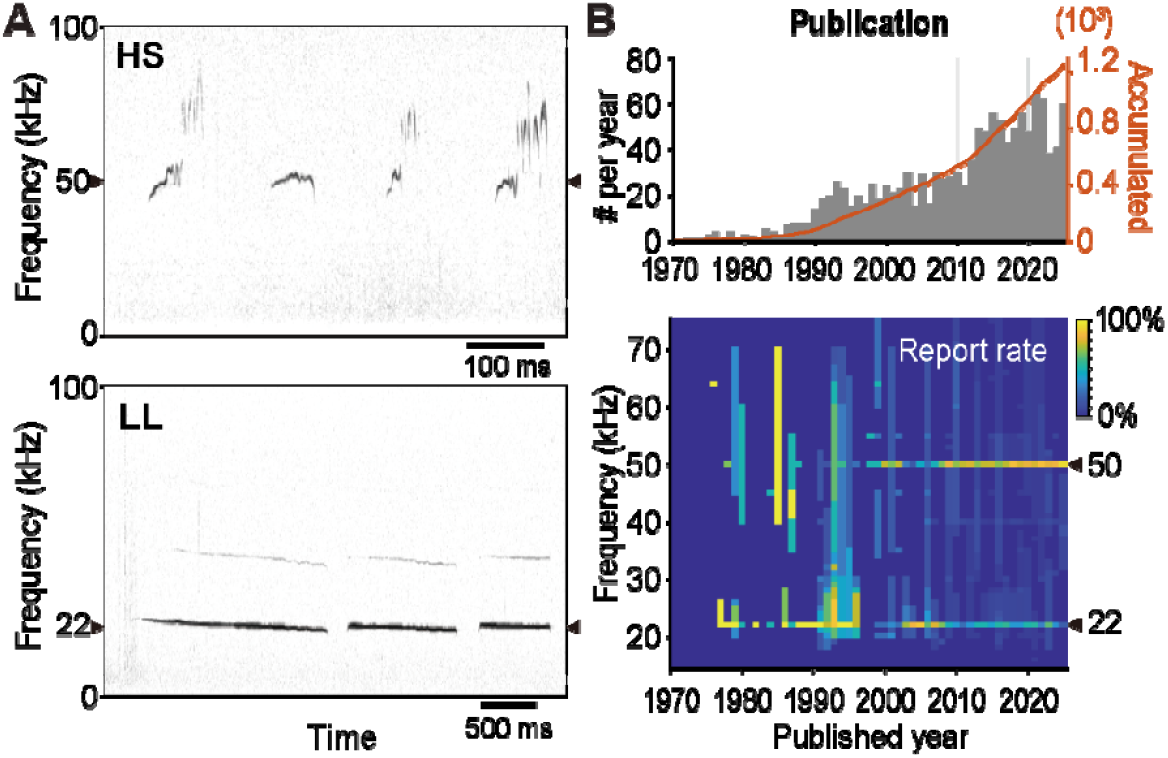
Recent publication on rat USVs. (*A*) Typical spectrograms of “high-short” (HS) and “low-long” (LL) calls in rat USVs. (*B*) The number of publications per year on rat USV (upper) and appearance ratio of frequency values (kHz) in the abstracts of publications per year (lower). Reports of 50 and 22 kHz have been predominant (as arrowheads point).

Over the past two to three decades, rat USVs have attracted considerable attention across multiple fields, including medical, pharmacological, and behavioral research, as evidenced by a substantial increase in related publications (***Fig. 1B, upper***). Examining the abstracts of papers, various frequency values were reported in studies until the early 1990s (***Fig. 1B, lower***). With the increase in publications after around 2000, most studies mentioned only two frequencies, 50 kHz or 22 kHz, not primarily to represent precise measured values but rather as convenient labels for vocalization categories. This suggests that a dichotomous (HS-vs-LL) classification of rat USVs has become increasingly dominant. Note that frequency ranges defining LL and HS calls are not consistent across research groups, leading to methodological variability. For example, frequency boundaries such as 30, 32, 33 kHz [17–20] have been used to distinguish LL calls (below the boundary) from HS calls (above the boundary). The widely adopted subtype scheme classified USVs spanning 20-95 kHz into LL calls (20-25 kHz) and 14 subtypes of HS calls [21].

Acoustic diversity within rat USVs has been documented as various subtypes within both HS and LL calls. For example, LL calls have been divided into long and short 22-kHz calls [22], while 14 subtypes have been identified within HS calls [21]. Some of these subtypes may have specific behavioral functions: for instance, flat 50-kHz calls may serve socio-coordinating functions [23], whereas frequency-modulated 50-kHz calls are more strongly connected to reward and positive affective states [24]. Although many studies have adopted such subtype schemes, most still begin by dividing USVs into the broader HS or LL categories before further subclassification. Thus, these subtypes are typically treated as subdivisions within the two main categories rather than as independent classes, and the dichotomous framework appears to remain prevalent.

However, several recent studies claim to have found new categories in rat USVs outside of the dichotomous framework along with evidence to infer their behavioral functions, *e*.*g*., 30 kHz for responding to HS calls [25]; 40 kHz for feeding [26]; 31 kHz for social inequality and isolation [27]. In light of these claims, we revisited the literature and found that several studies, including an early report more than a decade ago [28], had already described USVs whose acoustic features fall outside the dichotomous framework. Given these findings, we speculated that the practice of analyzing USVs by classifying them into HS or LL categories based on a single boundary might have undermined the possibility of detecting new vocalizations and corresponding behavioral contexts. Therefore, we systematically surveyed publications on rat USVs (***Fig. 2A***) to provide evidence supporting the existence of vocalizations outside the two conventional categories. We also present a perspective for future studies on the behavioral functions of USVs beyond the dichotomy.

**Figure 2.**
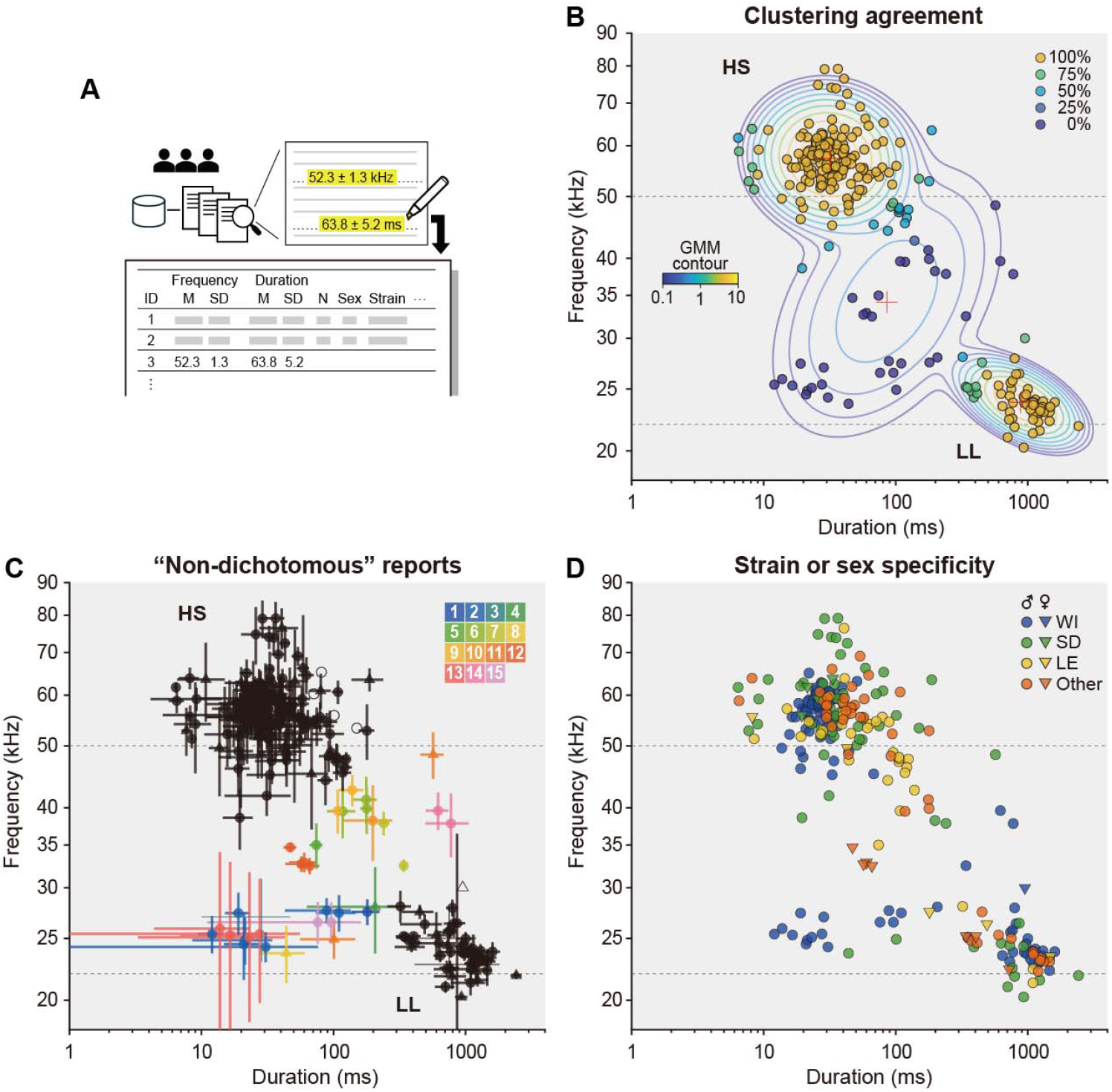
Distribution and clustering of collected data on a duration-frequency plane. (*A*) Procedures of the paper survey and extraction of numerical values. Over 400 articles were selected from those published before 2024, and the numerical values of frequency, duration, and other information in the text were extracted. (*B*) “Dichotomous”-likeliness of data and Gaussian mixture fitting. Filled circle shows representative frequency and duration reported in collected literature. Circle color indicates agreement rate (%) across four clustering methods for assigning the datapoint to the dichotomous call category (see ***Materials and Methods***). Contour line shows a mixture of three Gaussian distributions fitted to the dataset. Red cross indicates the center of distribution. Upper left and lower right clusters appear corresponding to HS and LL calls. (*C*) “Non-dichotomous” USVs identified by the clustering agreement (25% or less dichotomous likeliness). Colors other than black and numbers on the color legend correspond to respective articles reporting non-dichotomous USVs (see ***Table 1***). Thick and thin errorbars are s.d. and min-max ranges, respectively. Open circles show data that representative values were reported without ranges. Triangles indicate datapoints obtained from call-level but not animal-level samples. (*D*) Sex and strain difference showed no specificity in the acoustic features (WI: Wistar, SD: Sprague-Dawley, LE: Long-Evans).

## Results and Discussion

We found 417 articles based on an online database search and included 387 original research papers for rat USVs that were available in English. We then comprehensively examined the description of USV features reported there and eventually extracted 339 data points of USV frequency and/or duration from 74 articles (***Data S1***), excluding data obtained from pup calls or physiological/genetic manipulations (see Materials and Methods). We then selected 257 cases that contained both duration and frequency numbers for further analysis.

**Table 1.** Characteristics and literature information of non-dichotomous USVs.

| Plot ID | Reference | Frequency (kHz)<br>mean ± s.d. | Duration (ms)<br>mean ± s.d. | Sex | Strain |
| --- | --- | --- | --- | --- | --- |
| 1 | Haney and Miczek, 1993 [33] | 27.5 ± 2.5 | 180.0 ± 84.9 | F | LE |
| 2 | Schwartz et al., 2007 [20] | 24.8 ± 4.2 | 21.4 ± 25.7 | M | WI |
| | | 27.4 ± 4.1 | 19.0 ± 7.9 | M | WI |
| | | 25.4 ± 3.4 | 12.0 ± 5.2 | M | WI |
| | | 24.5 ± 5.9 | 20.9 ± 12.3 | M | WI |
| | | 27.6 ± 2.5 | 88.8 ± 91.7 | M | WI |
| | | 27.4 ± 3.6 | 110.0 ± 73.2 | M | WI |
| | | 24.2 ± 2.6 | 30.6 ± 91.2 | M | WI |
| 3 | Natusch and Schwartz, 2010 [34] | (31.0–23.0) | (47.0–10.0) | M | WI |
| 4 | Rao et al., 2014 [38] | 28.0 ± 8.6* | 206.5 ± 287.0* | both | WI |
| 5 | Burke et al., 2017 [32] | 35.0 ± 5.7 | 74.1 ± 16.3 | M | LE |
| 6 | Shimoju et al., 2020 [39] | 39.5 ± 7.3 | 117.6 ± 56.7 | M | WI/ST |
| | | 41.2 ± 7.1 | 176.6 ± 71.1 | M | WI/ST |
| | | 39.9 ± 7.0 | 178.5 ± 38.4 | M | WI/ST |
| 7 | Berz et al., 2022 [25] | 32.5 ± 1.5 | 340.0 ± 30.0 | M | WI |
| | | 37.8 ± 3.2 | 240.0 ± 50.0 | M | SD |
| 8 | Lara-Valderrábano et al., 2022 [36] | 23.7 ± 4.9 | 43.9 ± 33.1 | M | SD |
| 9 | Tryon et al., 2023 [40] | 42.6 ± 4.8 | 138.6 ± 58.5 | M | LE |
| | | 39.6 ± 6.3 | 107.8 ± 24.3 | M | LE |
| 10 | Cullins et al., 2023 [31] | 38.2 ± 10.4 | 198.0 ± 161.2 | M | SD |
| 11 | Bartsoen and Wöhr, 2023 [35] | 24.9 ± 3.4* | 100.8 ± 90.0* | M | SD |
| | | 48.4 ± 8.1* | 568.5 ± 229.0* | M | SD |
| 12 | Okabe et al., 2023 [27] | 32.9 ± 2.3 | 60.0 ± 27.7 | F | Lewis |
| | | 34.7 ± 1.1 | 47.0 ± 10.4 | F | Lewis |
| | | 32.7 ± 1.8 | 57.0 ± 27.7 | F | Lewis |
| | | 32.5 ± 1.9 | 66.0 ± 24.2 | F | Lewis |
| 13 | Olszyński et al., 2023 [29] | 25.9 ± 16.4 | 13.8 ± 18.9 | M | WI |
| | | 25.1 ± 13.2 | 23.1 ± 39.6 | M | WI |
| | | 25.4 ± 11.2 | 27.6 ± 56.3 | M | WI |
| | | 25.3 ± 15.3 | 16.5 ± 19.2 | M | WI |
| 14 | Olszyński et al., 2024 [30] | 37.8 ± 8.7 | 775.0 ± 559.5 | M | WI |
| | | 39.6 ± 5.4 | 619.6 ± 239.6 | M | WI |
| 15 | Perrier et al., 2024 [37] | 26.5 ± 4.0 | 96.0 ± 130.0 | M | WI |
| | | 26.5 ± 4.0 | 76.0 ± 130.0 | M | WI |

Abbreviations. M: male; F: female; SD: Sprague-Dawley; LE: Long-Evans; WI: Wistar ( ) Minimum to maximum range was reported.
\* Sample consisted of call counts (call-level) but not animals (animal-level).

Initially, we evaluated how these data clustered on a duration-frequency plane by fitting 2-D Gaussian distributions. Our analysis revealed that the data formed a mixture of two components corresponding to the dichotomous USVs (HS and LL calls) and an additional component encompassing other USVs (***Fig. 2B***). The centers of the three Gaussian components (C_1-3_) resembled the two representative frequency-duration profiles of the dichotomous USVs and a third set of values located between the typical HS and LL ranges (C_1_: 57.3 kHz, 31.7 ms; C_2_: 23.8 kHz, 880.3 ms; C_3_: 34.2 kHz, 85.6 ms; shown as red crosses in ***Fig. 2B***). We then applied four clustering methods (see Materials and Methods), which consistently identified distinct clusters around the two dichotomous USVs, suggesting that C_1_ and C_2_ corresponded to HS and LL calls, respectively. We classified data points as dichotomous USVs if at least two of the four clustering methods (≥50%) identified them as such. Although dichotomous USVs exhibited well-clustered distributions, 41 cases out of 257 data points fell outside the range of dichotomous USVs, which we termed “non-dichotomous” USVs (***Fig. 2C***; black and colored dots for dichotomous and non-dichotomous calls, respectively). Such cases were found in 15 articles [20,25,27,29–40] (***Table 1***). According to the result of the clustering agreement, data distributions on the duration-frequency plane exhibit sufficiently distinct clusters corresponding to dichotomous and non-dichotomous USVs, suggesting the existence of vocalizations outside the two conventional categories.

Our literature survey revealed that non-dichotomous USVs have been reported in 15 articles, which supports recent studies that claimed new categories of USVs outside the 50-/22-kHz dichotomous framework. Various types of non-dichotomous USVs have been repeatedly reported in both sexes and different strains (***Fig. 2D***), suggesting that such USVs are not exceptional or outliers nor biased toward any particular strain or sex, even though subtle acoustic differences have been previously reported [41,42]. Although causal manipulations of behavioral contexts are required to establish functional categories, the results indicate that there may be far more acoustic categories of USVs than previously thought.

Among papers reporting non-dichotomous USVs, some described those vocalizations beyond HS and LL calls as distinct categories, such as “short 22-kHz call” [29], “44-kHz call” [30], “30-kHz USVs” [31], “31-kHz call” [27], and “flat/trill combination call” [32]. Notably, the “short 22-kHz call” has been documented since a paper published about 30 years ago [22]. While its biological function remains unclear, it is thought to indicate internal discomfort [43]. Additionally, the “44-kHz call” has been noted in several recent publications and is suggested to be associated with aversive contexts [44,45]. Furthermore, some papers stated that the recorded vocalizations were atypical or differed in frequency and duration from the two main groups (HS or LL) [20,25,33–35]. Thus, among the fifteen papers reporting non-dichotomous USVs, ten papers explicitly mentioned their existence.

On the other hand, five papers documented non-dichotomous USVs but reported them as typical “22-kHz USVs” or “50-kHz USVs,” without explicitly mentioning non-dichotomous USVs [36–40]. For example, Shimoju et al. (2020) and Tryon et al. (2023) reported vocalizations with longer durations than typical HS calls as “50-kHz USVs”[39,40]. These vocalizations exhibited characteristics closer to “44-kHz calls” [30]. This discrepancy may arise because these studies focused on typical HS/LL USVs classified solely based on thresholds defined by a single frequency and a single duration. Therefore, even in papers that reported the existence of “50-kHz USVs” or “22-kHz USVs” but did not report the actual acoustic feature values of the vocalizations, the non-dichotomous USVs may have been observed in practice, although they were not included in the analysis.

Non-dichotomous USVs have been reported under various experimental conditions. Fear conditioning [29,30,40], tickling [20,34], and playback of conspecific vocalizations [25,29] were common experimental conditions across different studies. Additionally, less severe but somewhat unpleasant experimental conditions included: rats being threatened but protected from attack by an aggressive conspecific [33], rats accustomed to being stroked by the experimenter witnessing other rats being stroked instead [27], stroking stimulation of the ventral region [39], startle responses to noise [35], and sleep fragmentation [37]. Furthermore, non-dichotomous USVs were observed in situations related to social interactions with conspecifics, such as during face-to-face interactions [38], play fighting [32], and in males after mating [31]. Other experimental situations included when rats returned to their home cages after being removed [20] and after vehicle (0.9% NaCl) administration [36]. Among 15 articles reporting non-dichotomous USVs, the age of rats varied: approximately 7 weeks (juveniles to adolescents) in 4 studies, approximately 13 weeks (adolescents to adults) in 4 studies, 1 year (adults) in 1 study, a broad age range in 2 studies, and not specified in 3 studies (see ***Table S1*** for details). This indicates that non-dichotomous USVs are not restricted to a specific age class.

Our results do not negate the existence of traditional HS and LL USVs but suggest the existence of diverse USVs beyond the dichotomous framework. This raises the possibility that rat USVs can be mapped onto a continuum, not just a few categories. Supporting this idea, recent studies revealed that laboratory mouse USVs do not show clear clusters but form a continuum [46,47]. Although this hypothesis cannot be tested with the current dataset, future analyses using larger, openly available datasets could address this question. Making raw datasets of rat USVs publicly available, together with clear descriptions of acoustic parameters and analysis procedures, would greatly facilitate such analyses.

Our approach had several limitations. First, because our dataset was extracted from published articles that used heterogeneous and sometimes insufficiently documented analysis procedures, the acoustic measurements reported may not always be fully reliable. Second, limitations also arise from the reporting formats and preprocessing methods (e.g., threshold settings for call detection or noise reduction) adopted in the surveyed papers. In this study, we extracted as much information as possible regarding the frequency and duration of USVs reported in the papers. Therefore, for example, multiple data points obtained from the same paper were treated as separate data points unless there was data redundancy. This may have contributed to the clustering effect. Furthermore, the values extracted from the papers and used for the clustering analysis were the average frequencies and durations. Considering that rat USVs were often divided into and then averaged within groups of HS or LL USVs, it was possible that some vocalizations, despite actually being non-dichotomous USVs, were forced into the traditional HS/LL USV groups during the averaging process. Finally, the analysis was based solely on frequency and duration. While focusing on frequency and duration is a standard approach— spectrograms (with time on the horizontal axis and frequency on the vertical axis) are typically examined first when analyzing animal vocalizations—the apparent existence of non-dichotomous USV groups might stem from relying solely on frequency/duration analysis. Since acoustic features exist beyond these two parameters, future studies employing analyses based on other acoustic characteristics (*e*.*g*., sound pressure or frequency modulation) would be interesting.

In conclusion, our study supports the abundance of non-dichotomous USVs beyond the dichotomous framework, emphasizing the importance of research on diverse vocalizations. We encourage future studies to report original acoustic measurements and analysis procedures, rather than relying solely on the simple distinction between HS and LL calls. Approaching rat USVs from a non-dichotomous perspective will provide deeper insights into the mechanisms of emotional communication in animals.

## Materials and Methods

### Paper survey and data collection

We searched for original papers related to rat USVs in the literature database (“Web of Science”). We collected numerical descriptions of USV data (duration, frequency, sample size, sex, and strain) in papers. For the literature survey, we used the following search key in the Web of Science: “AB=(rat (ultrasonic OR kHz) (call OR vocalization) NOT(pup OR infant OR neonatal))”, Document type: “Article”, Citation Topics Micro: “1.5.1511 Ultrasonic Vocalization”. We only included papers that were classified in the following “WoS Categories”: *Neurosciences, Behavioral Sciences, Pharmacology Pharmacy, Psychiatry, Psychology Biological, Zoology, Multidisciplinary Sciences, Clinical Neurology, Veterinary Sciences, Psychology, Psychology Multidisciplinary, Biology, Audiology Speech Language Pathology, Developmental Biology, Endocrinology Metabolism, Substance Abuse, Medicine Research Experimental, Acoustics, Genetics Heredity, Physiology, Anesthesiology, Linguistics, Cell Biology, Ecology, Engineering Biomedical, Environmental Sciences, Evolutionary Biology, Otorhinolaryngology, Psychology Experimental*. As a result, we found 398 articles (searched on April 18, 2024). We further added 19 papers that authors found by chance during the article survey or revision processes, and hence, the literature list size became 417. After collecting these articles, we started to read the literature and omitted articles that were reviews or book chapters but not original papers.

To extract the USV data (duration, frequency, sample size, sex, and strain), we draw highlights on the corresponding descriptions in paper PDFs using the reference management software Zotero. We excluded USV data obtained from pups, while retaining adolescent and juvenile calls. We also excluded data from electrophysiological or optogenetic stimulation experiments and from genetically or pharmacologically manipulated animals, whereas wild-type and saline-treated control animals were included.

Many papers in our collection described the “mean frequency” or “peak frequency” as the frequency characteristic of USVs. The mean frequency typically indicates the mean of the frequency trace in a call, whereas the peak frequency is the frequency at the time point where the amplitude showed its peak in the call. In rare cases, we found different descriptions, such as “representative frequency” or “dominant frequency.” We labeled these characteristics differently during the data extraction. The USV characteristics were typically described as “mean ± standard deviation (s.d.)” or “mean ± standard error (s.e.)” in most articles that were calculated across subjected animals. In some cases, we also found median values or the value range as “minimum - maximum” forms. We labeled these statistics differently during the data extraction. Note that we collected data with the frequency at ≥19 kHz to include only ultrasonic calls.

In many studies, summary statistics such as mean, s.d., and min.-max. range were calculated at the individual level based on per-animal representative values of call features; but some studies computed directly from all call-level feature values, without first aggregating within an animal (either from all calls produced by a single animal or from calls pooled across multiple animals). We chose the former (statistics of animal-level samples) instead of the latter (call-level) if the article reported both. Additionally, we converted the s.e. value to s.d. if we could obtain the corresponding sample size (*n*) by multiplying the s.e. value by the square root of *n*.

### Distribution assessment

To explore the presence of calls beyond the dichotomous USVs (HS and LL calls), we examined the distribution of the collected data points on a duration-frequency plane. In this analysis, we included only those data points that had both duration and frequency values, regardless of the type of frequency characteristics (such as mean frequency, peak frequency, or others) for simplification. We treated different statistical measures (mean, median, and min-max range) equally, disregarding differences at the sample level (whether individual- or call-level). For range data, we calculated the midpoint between the minimum and maximum values to serve as the representative number. We log-transformed duration and frequency to stabilize variance across the range of values; otherwise, variability on the linear scale would increase with larger duration and frequency and bias the assessment.

We started by determining the optimal number of clusters by fitting a Gaussian mixture model (GMM), which is a weighted sum of multiple Gaussian distribution components, while varying the number of components. By repeatedly calculating the Bayesian information criterion (BIC) to evaluate the goodness of fit, we consistently found that the ideal number of components ranged between four and six. This finding indicates that the collected data consist of more than just two dichotomous clusters. We then approximated the data using GMM with three components for visualization purposes. The centers of the fitted three components resembled the two representative frequencies of the dichotomous USVs and a frequency value between them.

Next, we categorized data points into three groups (HS, LL, and other calls) using four clustering algorithms (GMM-based clustering, DBSCAN [48,49], OPTICS [50], and BIRCH [51]), implemented in MATLAB (Statistics and Machine Learning Toolbox, R2024b) and Python (scikit-learn library, version 1.5.2, Python version 3.12):

1. Clustering based on the previously obtained Gaussian mixture approximation.
2. DBSCAN, which identifies clusters as high-density areas separated by low-density regions (parameter setting: *eps* = 0.4, *min_samples* = 20).
3. OPTICS, similar to DBSCAN but with relaxed constraints (*min_samples* = 40, *max_eps* = 0.6).
4. BIRCH, which approximates a hierarchical, tree-like structure of subclusters (*threshold* = 0.4, *branching_factor* = 30, *n_clusters* = 4).

We standardized the frequency and duration values by dividing them by their standard deviation after logarithmic conversion. For the first three methods, parameters were optimized to clearly visualize clusters corresponding to the dichotomous USVs (HS and LL) while dividing data into three clusters. For BIRCH, the maximum number of clusters was set to be four to help this method identifying the two core regions corresponding to the dichotomous USVs.

All four clustering methods consistently detected two core regions corresponding to high-short and low-long calls. Additionally, each method identified a third cluster separate from these core regions (classified as outliers by DBSCAN and OPTICS, and as two combined clusters by BIRCH), suggesting the existence of non-dichotomous USVs.

### Yearly publication and frequency reporting rates in paper abstracts

To investigate the change over time in the number of papers published that mention rat vocalization, we searched the following search key in the Pubmed paper database: “((vocalization [Title/Abstract]) OR (vocalizations [Title/Abstract]) OR (vocalisation [Title/Abstract]) OR (vocalisations [Title/Abstract])) AND (ultrasonic [Title/Abstract]) AND ((rat [Title/Abstract]) OR (rats [Title/Abstract]))”. As a result, 1266 papers were found (searched on December 5, 2025). The number of published papers was counted per year.

To assess yearly changes of USV frequency reports in paper abstracts, we searched for USV-related papers with the following search key in the Web of Science: “AB=(kHz rat (vocalization OR call))”. As a result, 599 papers were found (searched on December 5, 2025). We searched for frequency descriptions using pattern matching (*e*.*g*., “X kHz”, “X-Y kHz”, “between X and Y kHz”). Unclear range descriptions (*i*.*e*., shown with “above”, “higher than”, “>“, “<“, etc.) were excluded from further analysis since almost all of them were related to analysis or stimulus conditions. After these extractions, we counted the detection for every frequency number from 15 to 80 kHz in 1 kHz steps. When the same frequency was reported multiple times in one paper, we counted it only once to avoid redundancy. For the range expressions, we added one for all frequency bins within that range. We then created a two-dimensional histogram with publication year as the horizontal axis and frequency as the vertical axis. The raw histogram was divided by the number of papers that reported the frequency in their abstracts to obtain the frequency reporting rates for each publication year.

## Author Contributions

RW, SH, and ROT wrote the manuscript. ROT performed data processing, visualization, and preparation of supplementary information. RW, SH, RK, and GT initiated the USV data survey project. All authors participated in the survey. SO and KO supervised the study. All authors were involved in revising and editing the manuscript.

## Competing Interest Statement

The authors declare no competing interest.

## Supporting Information

**Data S1**. All collected data of duration, frequency and other information (74 papers).

**Table S1.**
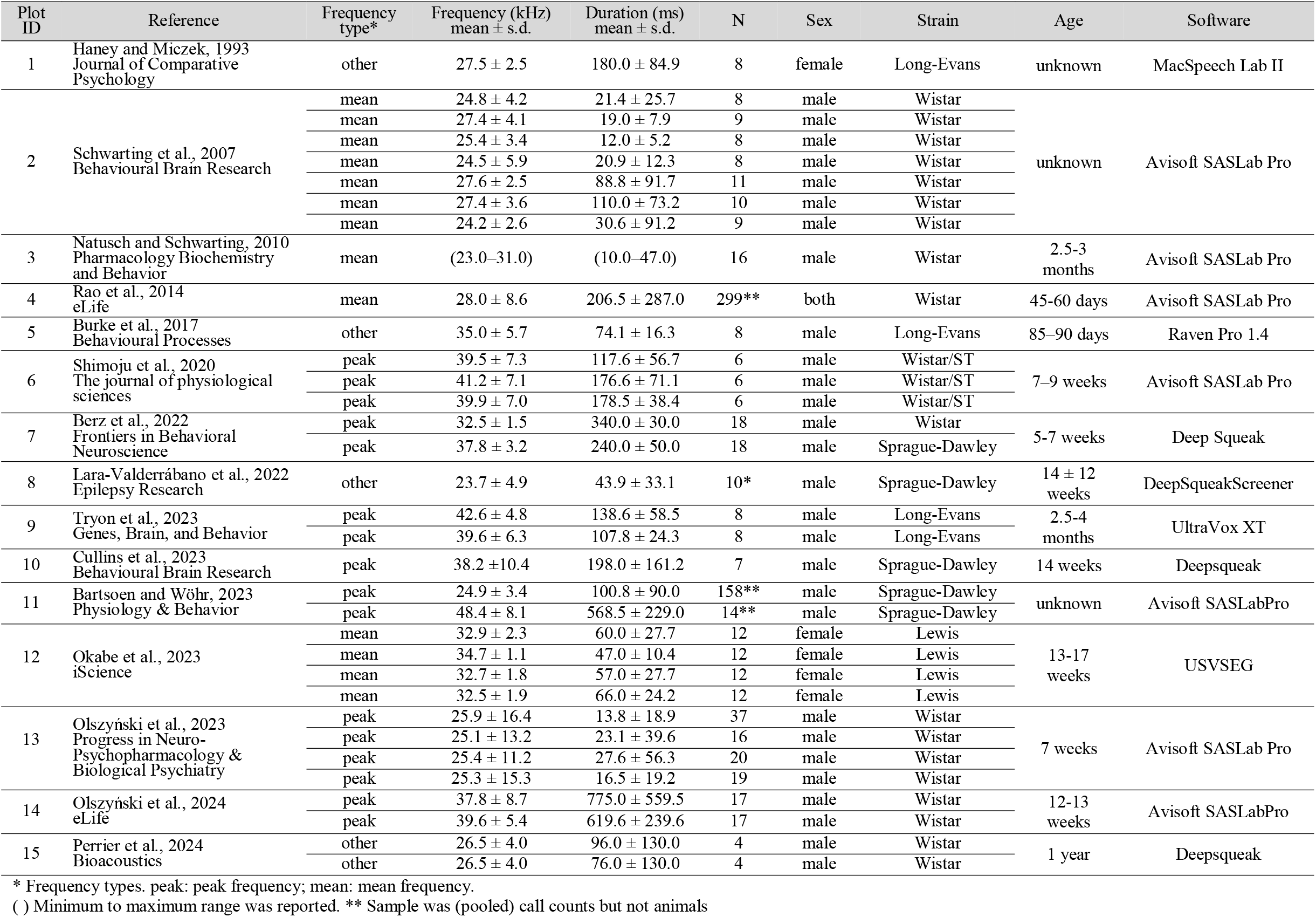
Detailed characteristics and additional information for non-dichotomous USVs.

## Notes

### Competing Interest Statement

The authors have declared no competing interest.

### Summary of Updates

All sections have been updated. We have added numerous references and expanded the introduction and discussion sections. Data sets have also been updated.

### Funding

- Japan Society for the Promotion of Science: 23H05428, 23H03243, 23H04342, 22H05157, 24K06624, 23H04336
- Japan Society for the Promotion of Science: 22KJ0702, 25KJ0778
- Japan Science and Technology Agency: JPMJPR22S8, JPMJFR2151, JPMJFR2320
- Smoking Research Foundation

